# Activity in developing prefrontal cortex is shaped by sleep and sensory experience

**DOI:** 10.1101/2022.07.31.502200

**Authors:** Lex J. Gómez, James C. Dooley, Mark S. Blumberg

## Abstract

In developing rats, behavioral state exerts a profound modulatory influence on neural activity throughout the sensorimotor system, including primary motor cortex (M1). We hypothesized that similar state-dependent modulation occurs in higher-order cortical areas with which M1 forms functional connections. Here, using 8- and 12-day-old rats cycling freely between sleep and wake, we record neural activity in M1, secondary motor cortex (M2), and medial prefrontal cortex (mPFC). At both ages in all three areas, neural activity increased during active sleep (AS) compared with wake. Regardless of behavioral state, neural activity in M2 and mPFC increased during periods when limbs were moving. This movement-related activity in M2 and mPFC, like that in M1, is driven by sensory feedback. Our results, which diverge from those of previous studies using anesthetized pups, demonstrate that AS-dependent modulation and sensory responsivity extend to prefrontal cortex. These findings expand the range of factors shaping the activity-dependent development of higher-order cortical areas.

## INTRODUCTION

The functional development of cerebral cortex is a sinuous and often surprising process, even for those structures with the most transparent of adult functions. Consider primary motor cortex (M1), whose name reflects its well-established role in adult motor control: In early development, M1 does not contribute to motor control at all, but instead functions exclusively as a sensory structure (Bruce et al., 1980; Chakrabarty and Martin, 2005; Young et al., 2012; Tiriac et al., 2014; Dooley and Blumberg, 2018; Singleton et al., 2021). The early-emerging somatosensory map in M1 provides the foundation upon which its later-emerging motor map is built (Dooley and Blumberg, 2018).

Another surprising aspect of M1 in early development is that its activity is modulated by behavioral state, in particular active sleep (AS, or REM sleep). In infant rats, this modulation reflects AS-dependent increases in neural activity that are enhanced by limb movements during AS, called twitches, that discretely and preferentially trigger sensory feedback to M1 and other sensorimotor structures (Blumberg, 2015; Blumberg et al., 2020). Given that AS predominates in early life (Jouvet-Mounier et al., 1969; Gramsbergen et al., 1970), it has been posited that this sleep state plays an outsized role in typical and atypical development (Blumberg et al., 2020).

That sleep so profoundly modulates neural activity in developing M1 raises the possibility that it also modulates higher-order prefrontal cortical areas with which M1 forms reciprocal connections, such as secondary motor cortex (M2) and medial prefrontal cortex (mPFC) (van Eden et al., 1992; Bedwell et al., 2014). As its name implies, M2 has a particularly close functional and anatomical connection with M1, integrating multimodal sensory cues for motor planning and modulating M1 activity during goal-directed action (Yin, 2009; Sul et al. 2011; Omlor, 2016; Barthas and Kwan, 2017; Morandell and Huber, 2017; Wang et al., 2020). Like M1, M2 develops a somatotopic map, further highlighting its dependence on sensory input (Yin, 2009; Kunori and Takashima, 2016; Omlor, 2016; Barthas and Kwan, 2017; Chen et al., 2017; Singleton et al., 2021). Thus, we hypothesize that, in infant rats, M2 is similar to M1 in that it responds to sensory input and is modulated by behavioral state.

In contrast with M1 and M2, mPFC in adults is not closely associated with sensorimotor functions, but rather with cognitive processes such as decision-making and attention (Tanji and Hoshi, 2001, 2008; Miller et al., 2002; Barbas and Zikopoulos, 2007; Euston et al., 2012). In infant rats, it is not known whether behavioral state modulates activity in mPFC, nor is it known whether mPFC processes sensory input. In fact, it has been theorized that prefrontal cortex, including mPFC, develops its unique higher-order functions precisely because—in contrast with M1 and other sensory-dependent cortical areas—it develops independently of sensory input (Johnson et al. 2015).

What is currently known about functional development in mPFC derives primarily from neural recordings from rat pups under urethane anesthesia (Brockmann et al., 2011; Bitzenhofer et al., 2015). Urethane precludes natural sleep-wake cycles, but allows expression of spindle bursts in mPFC (Brockmann et al., 2011). Spindle bursts are brief thalamocortical oscillations that, in primary sensory areas, are closely associated with the processing of sensory stimuli (Khazipov et al., 2004; Hanganu et al., 2007; Dooley et al., 2020). In the mPFC of urethanized pups, however, spindle bursts appear to occur spontaneously. Here, we determine if this is also the case in unanesthetized pups, as well as test the hypothesis that the infant mPFC, like M1, is modulated by behavioral state.

Using unanesthetized rats at postnatal days (P) 8 and P12, we find that M2 and mPFC exhibit state-dependent modulation such that neural firing rates are highest during AS, especially during periods of twitching. We also find that neurons in M2 and mPFC respond to sensory input arising from limb movements that are self-generated (i.e., reafference) or other-generated (i.e, exafference). Finally, to explain discrepancies between the present findings and those reported earlier, we show that urethane administration at P8 prevents expression of behavioral state and brain-behavior relations. Altogether, these findings demonstrate that the previously documented effects of behavioral state and sensory experience on somatosensory activity in M1 extend to M2 and mPFC, thus pointing to new directions for conceptualizing activity-dependent development of higher-order cortical areas.

## RESULTS

We recorded extracellular unit activity in M1, M2, and mPFC in head-fixed rats at P8 and P12 (**Fig. 1A**). For each pup, recordings were performed first in the forelimb regions of M1 and M2 for 40 min, followed by 50 manual stimulations of the forelimb contralateral to the recording sites; this procedure was repeated for recordings in M1 and mPFC. Electrode locations in M1, M2, and mPFC were confirmed histologically (**Fig. 1B**). At P8, we collected eight M1-M2 recordings (107 M1 units, 118 M2 units) and eight M1-mPFC recordings (117 M1 units; 103 mPFC units); at P12 we collected nine M1-M2 recordings (217 M1 units; 204 M2 units) and eight M1-mPFC recordings (222 M1 units; 179 mPFC units). Neural activity, electromyographic (EMG) activity in the nuchal and biceps muscles, and high-speed video (100 frames/s) were recorded as pups cycled between sleep and wake (**Fig. 1C-D**). As expected (Dooley et al., 2021; Glanz et al., 2021; Gómez et al., 2021), pups spent more time in AS than wake at P8 (AS: 57.7 ± 2.5%; wake: 30.9 ± 1.9%) and P12 (AS: 44.0 ± 3.6%; wake: 39.3 ± 3.5%). Also, the transition from discontinuous cortical activity at P8 to continuous activity at P12 is evident in all three areas, as described previously in primary somatosensory, motor, and visual cortex (Golshani et al., 2009; Rochefort et al., 2009; Van Der Bourg et al., 2017; Glanz et al., 2021; Riyahi et al., 2021).

**Figure 1.**
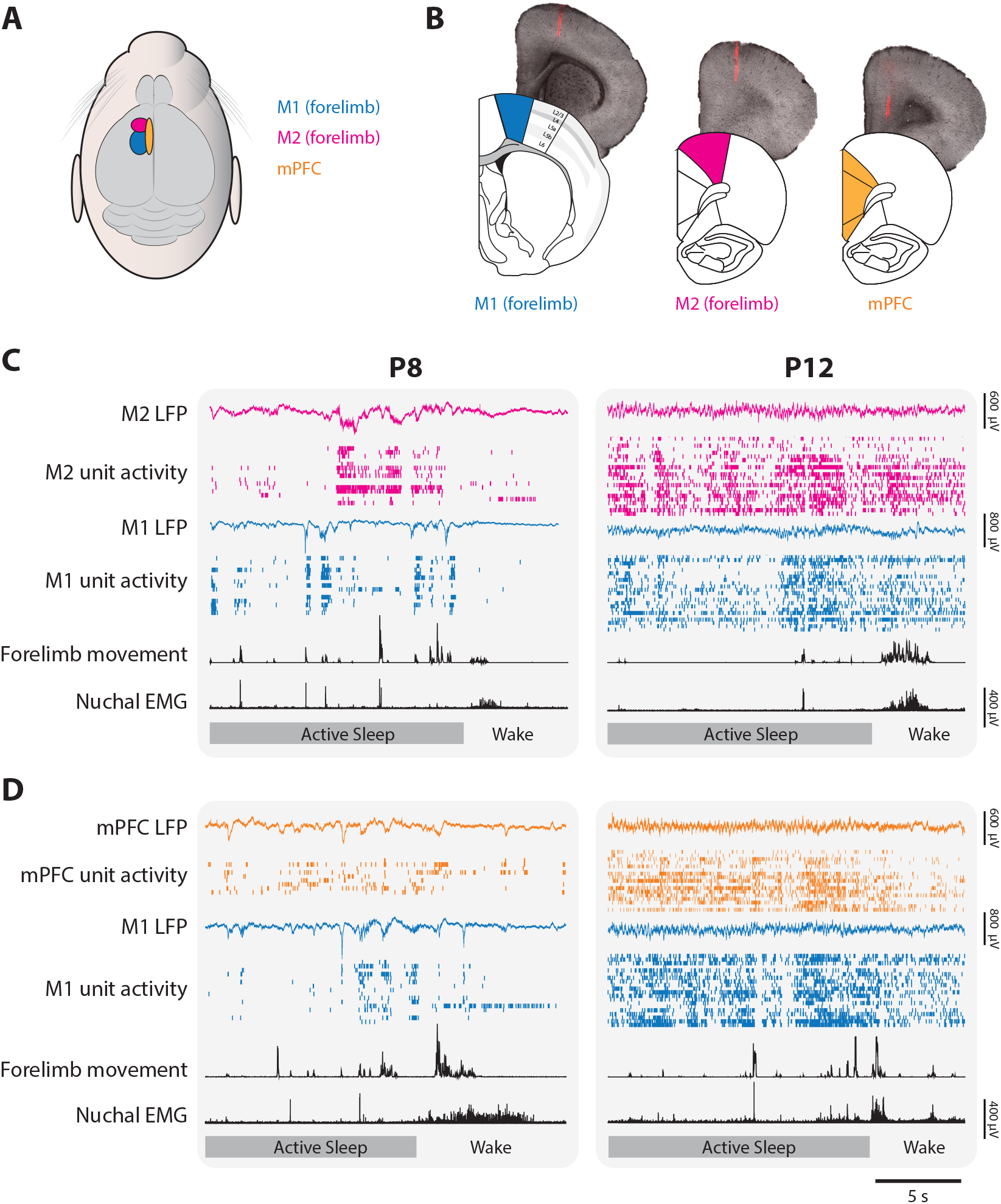
Representative neural activity in M1, M2, and mPFC in P8 and P12 rats. (A) Illustration showing the surface locations of M1 (blue), M2 (magenta), and mPFC (gold). These color codes are used in all other figures. (B) From left to right, illustrations of coronal sections of M1, M2, and mPFC below corresponding brightfield coronal sections showing a fluorescent electrode track in each area. (C) Representative 20-s segments of data from recordings in M1 and M2 at P8 (left) and P12 (right) across behavioral states. For each record from the top, data are presented as follows: M2 LFP (magenta trace), M2 unit activity (magenta ticks), M1 LFP (blue trace), M1 unit activity (blue ticks), forelimb movement, and nuchal EMG. Bottom row: Behavioral states marked as active sleep or wake. (D) Same as in C, but with mPFC instead of M2 (mPFC LFP, gold trace; mPFC unit activity, gold ticks).

### Neural activity in M2 and mPFC is modulated by behavioral state

At P8, representative recordings in M1, M2, and mPFC illustrate substantial and often abrupt increases in neural activity during AS (**Fig. 2A**). In each area, the mean firing rate was significantly higher during AS than wake (t_(7)_s ≥ 4.38, ps ≤ 0.003, Cohen’s Ds ≥ 1.55; **Fig. 2B**). State-dependent modulation of cortical activity continued through P12 (**Fig. 2C**); once again, the mean firing rate in each area was significantly higher during AS than wake (t_(7-8)_s ≥ 3.17, ps ≤ 0.016, Cohen’s Ds ≥ 1.12, **Fig. 2D**). Similarly, at P8, the rate of spindle bursts was higher during AS than wake for all three areas (t_(7)_s ≥ 3.805, ps ≤ 0.007, Cohen’s Ds ≥ 1.35; **Fig. 3**). (Spindle bursts were not analyzed at P12 due to their low rate of occurrence at this age.) Thus, like M1, neural activity in M2 and mPFC is modulated at both ages in a state-dependent manner.

**Figure 2.**
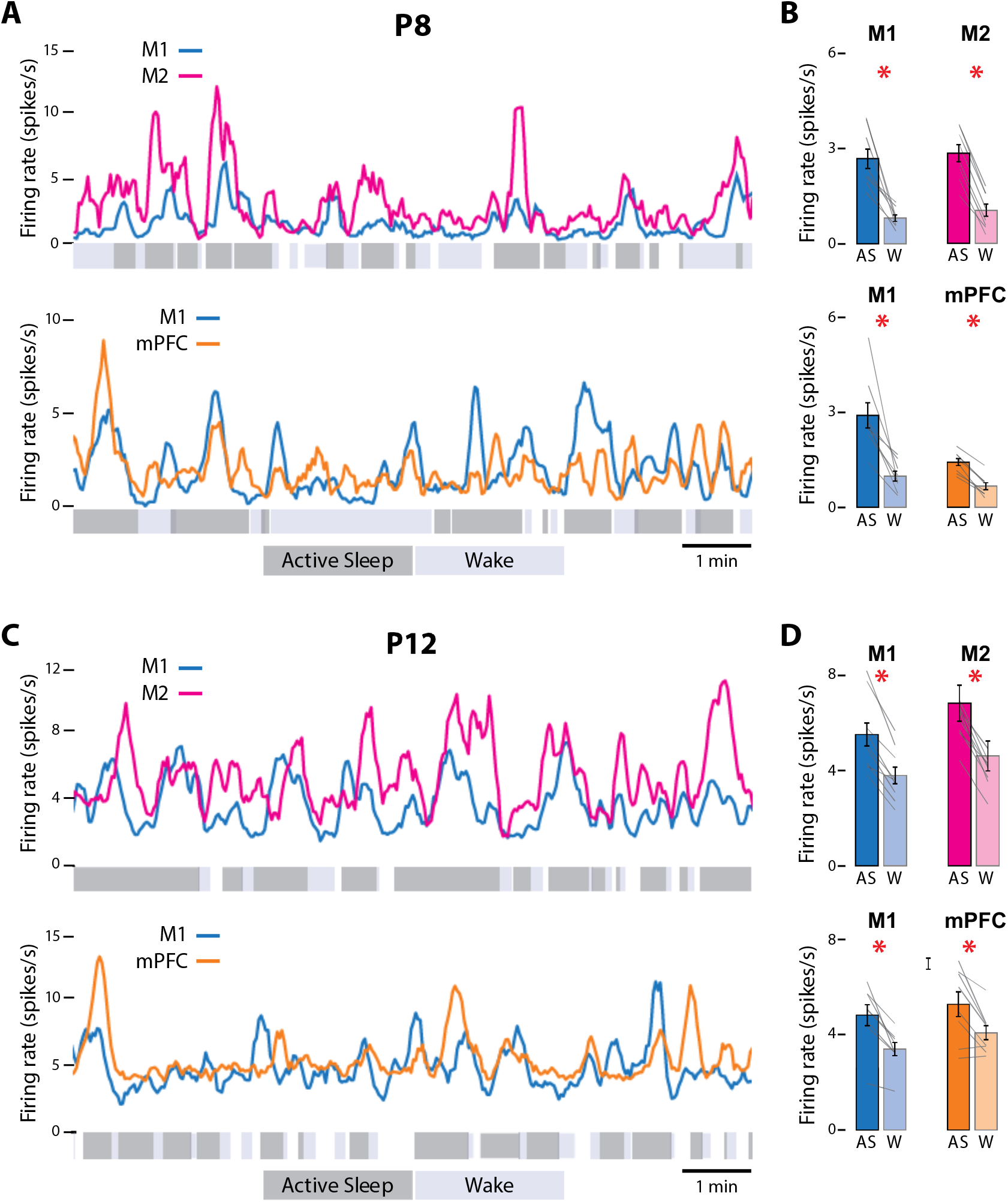
State-dependent unit activity in M1, M2, and mPFC in P8 and P12 rats. (A) Representative 10-min segments of data from a P8 rat showing mean firing rate (2-s bins) in relation to active sleep (dark grey) and wake (light grey). Top: Units in M1 and M2. Bottom: Units in M1 and mPFC. (B) Top: Mean firing rates for M1 and M2 units during active sleep (AS) and wake (W). Bottom: Mean firing rates for M1 and mPFC units during AS and wake. Mean firing rates for individual pups are shown as grey lines. Error bars are SEM. Asterisks denote significant difference between states, p ≤ 0.025. (C) Same as in A, but for a P12 rat. (D) Same as in B, but for P12 rats. (For M2, the values for one data pair exceed 8 spikes/s and are not shown.)

**Figure 3.**
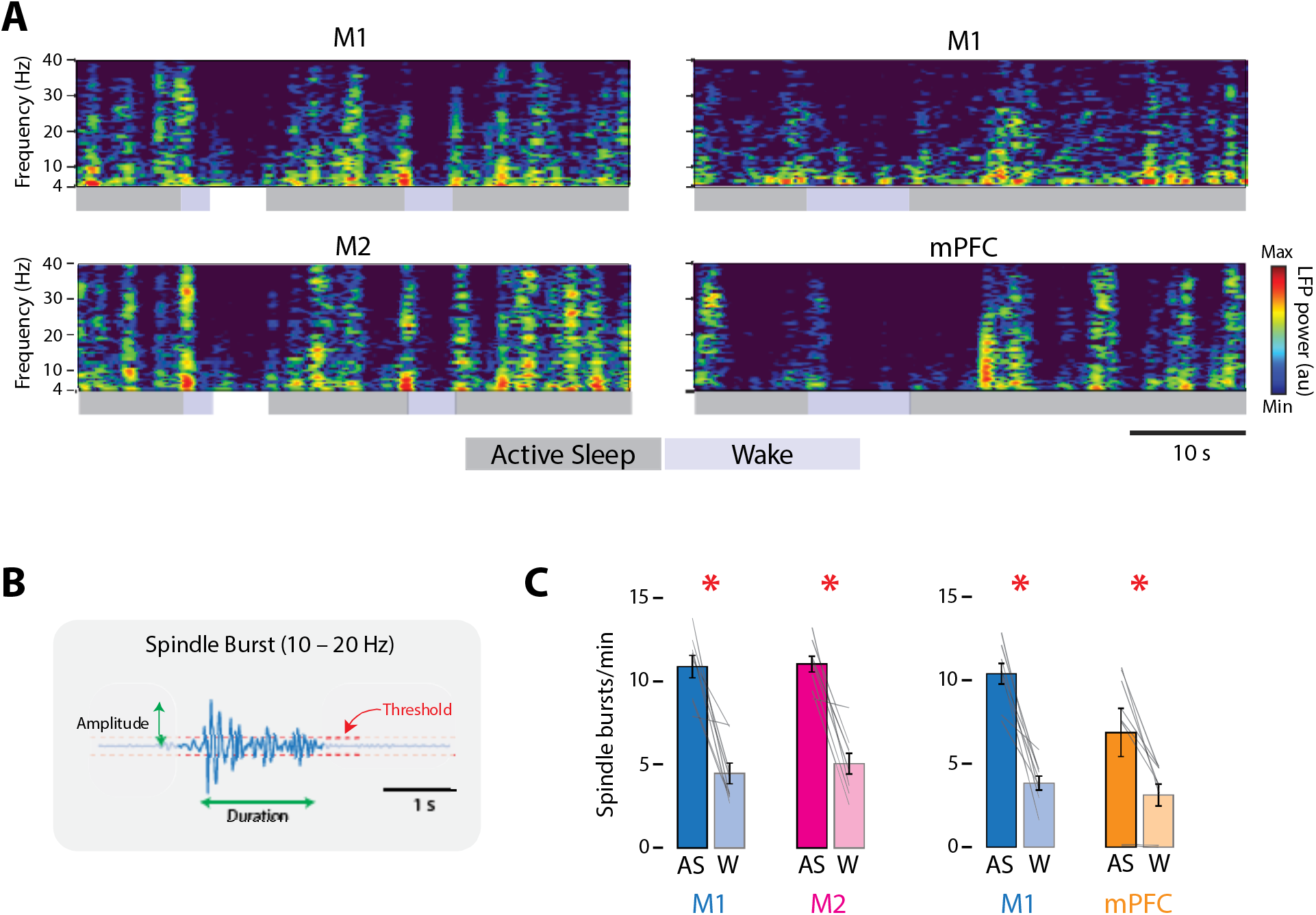
State-dependent LFP activity in M1, M2, and mPFC in P8 rats. (A) Left column: Representative 50-s segment of data showing spectrogram for M1 (top) and M2 (bottom) across active sleep (dark grey) and wake (light grey). Right column: Same as for left column, but for M1 (top) and mPFC (bottom). (B) Illustration to show method for detecting spindle bursts and calculating their amplitude and duration. (C) Bar graphs showing mean spindle-burst rate in M1, M2, and mPFC during active sleep (AS) and wake (W). Mean firing rates for individual pups are shown as grey lines. Error bars are SEM. Asterisks denote significant difference between states, p ≤ 0.025.

### Neural activity in M2 and mPFC increases during periods of self-generated movement

In infant rats, AS-dependent increases in M1 activity correspond with periods of limb movement (e.g., Glanz et al. 2021). We next determined whether the same is true for M2 and mPFC (**Fig. 4A**). At both ages, the mean firing rate in each area increased significantly during periods of movement (**Fig. 4B**). For all cases, repeated-measures ANOVAs revealed significant main effects of behavioral state (F_(1,7-8)_s ≥ 16.90, ps ≤ 0.005, η_p_^2^s ≥ 0.71) and movement (F_(1,7-8)_s ≥ 6.83, ps ≤ 0.035, η_p_^2^s ≥ 0.49). None of the state x movement interactions was significant (F_(1,7-8)_s ≤ 5.35), except for one of the M1 tests at P8 (F_(1,7)_ = 6.11, p = 0.043, η_p_^2^ = 0.47).

**Figure 4.**
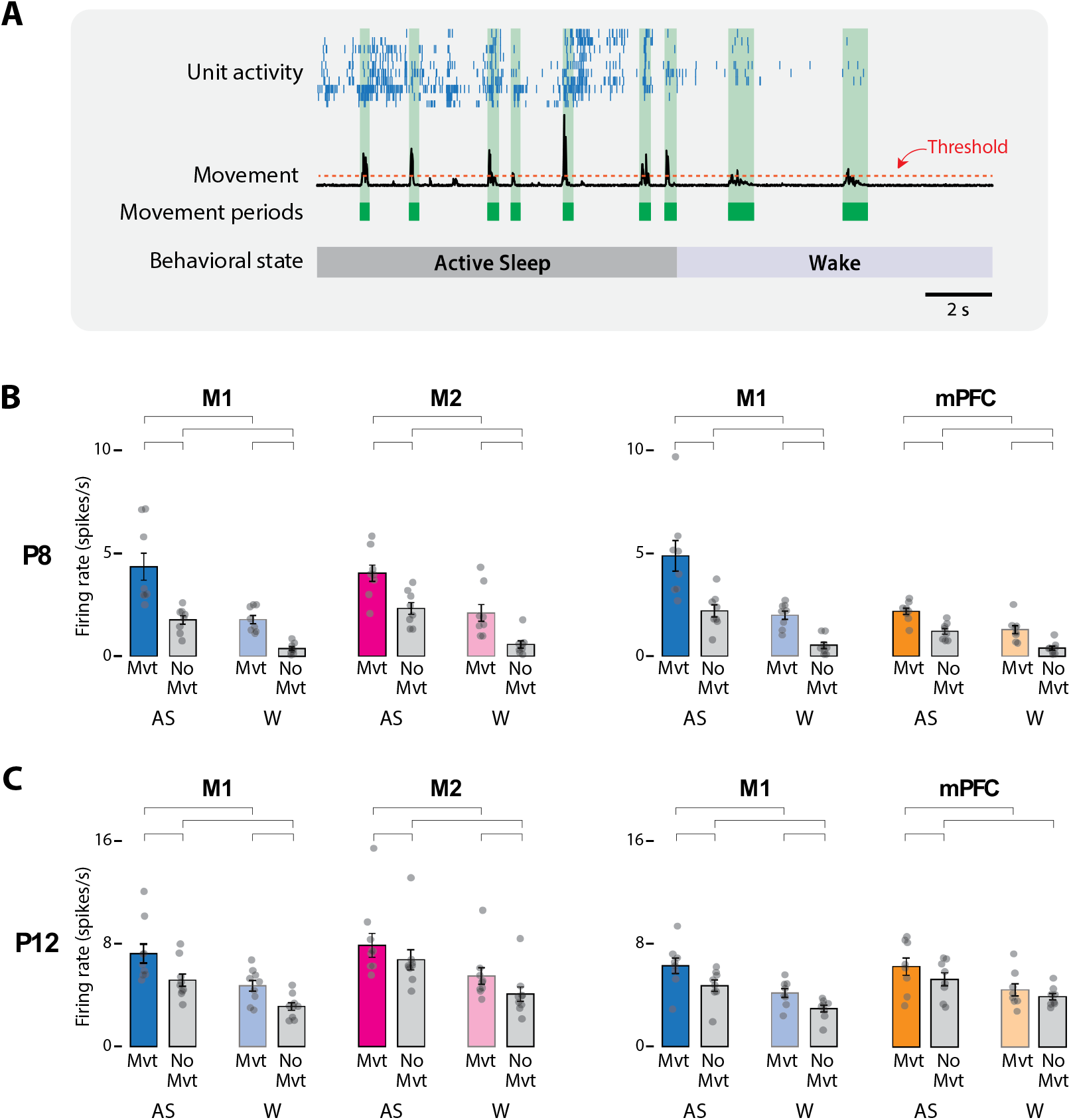
Movement-dependent unit activity in M1, M2 and mPFC in P8 and P12 rats. (A) Representative 20-s segment of data showing unit activity (blue ticks), movement data (black trace), movement periods (green blocks), movement-detection threshold (dotted orange line), and behavioral state. (B) Bar graphs showing mean firing rates for neurons in M1, M2, and mPFC during periods of movement (Mvt) or no movement (No Mvt) across AS and wake (W). Mean firing rates for individual pups are shown as grey circles. Error bars are SEM. Brackets denote significant difference between groups, p ≤ 0.0125. (C) Same as in B, but for P12.

For each of the eight repeated-measures ANOVAs across P8 and P12, four planned comparisons were conducted to compare firing rates within behavioral state across movement conditions and within movement conditions across behavioral state (**Fig. 4B-C**). Of the 32 planned comparisons, 31 were significant (t_(7-8)_s ≥ 3.51, ps ≤ 0.01, Cohen’s Ds ≥ 1.24). The general pattern was for firing rates to be highest during AS-related periods of movement (i.e., twitching), intermediate during periods of AS-related periods of quiescence and wake-related periods of movement, and lowest during wake-related periods of quiescence.

In summary, at P8 and P12, neural activity in M1, M2, and mPFC reflects the interactive effects of behavioral state and movement. Given that all three areas exhibited similar movement-related increases in activity and that M1 is known to respond to movement-related sensory feedback (Dooley and Blumberg, 2018; Gómez et al., 2021), we determined next whether M2 and mPFC are also responsive to sensory input.

### Neurons in M2 and mPFC respond to sensory input

We quantified neural responses in M1, M2, and mPFC to forelimb twitches, wake movements, and stimulations (**Fig. 5A**). As expected, at both ages, units in M1 and M2 responded to sensory feedback from twitches, wake movements, and stimulations; surprisingly, so did units in mPFC. The percentage of responsive units in all three areas varied by age and event type. In general, at P8, M1 exhibited the highest percentage of responsive units, followed by M2 and then mPFC (t_(7)_s ≥ 4.44, ps ≤ 0.003, Cohen’s Ds ≥ 1.57; **Fig. 5B**). Mean M2 responsiveness was 60.9 ± 10.2% for twitches, 46.5 ± 11.9% for wake movements, and 36.7 ± 9.1% for stimulations; for mPFC, these values were 37.8 ± 8.0%, 20.6 ± 7.5%, and 9.4 ± 5.5%, respectively. At P12, responsiveness declined to low levels in all three areas, but M1 was still more responsive than M2 or mPFC (t_(7-8)_s ≥ 3.13, ps ≤ 0.017, Cohen’s Ds ≥ 1.11; **Fig. 5C**).

**Figure 5.**
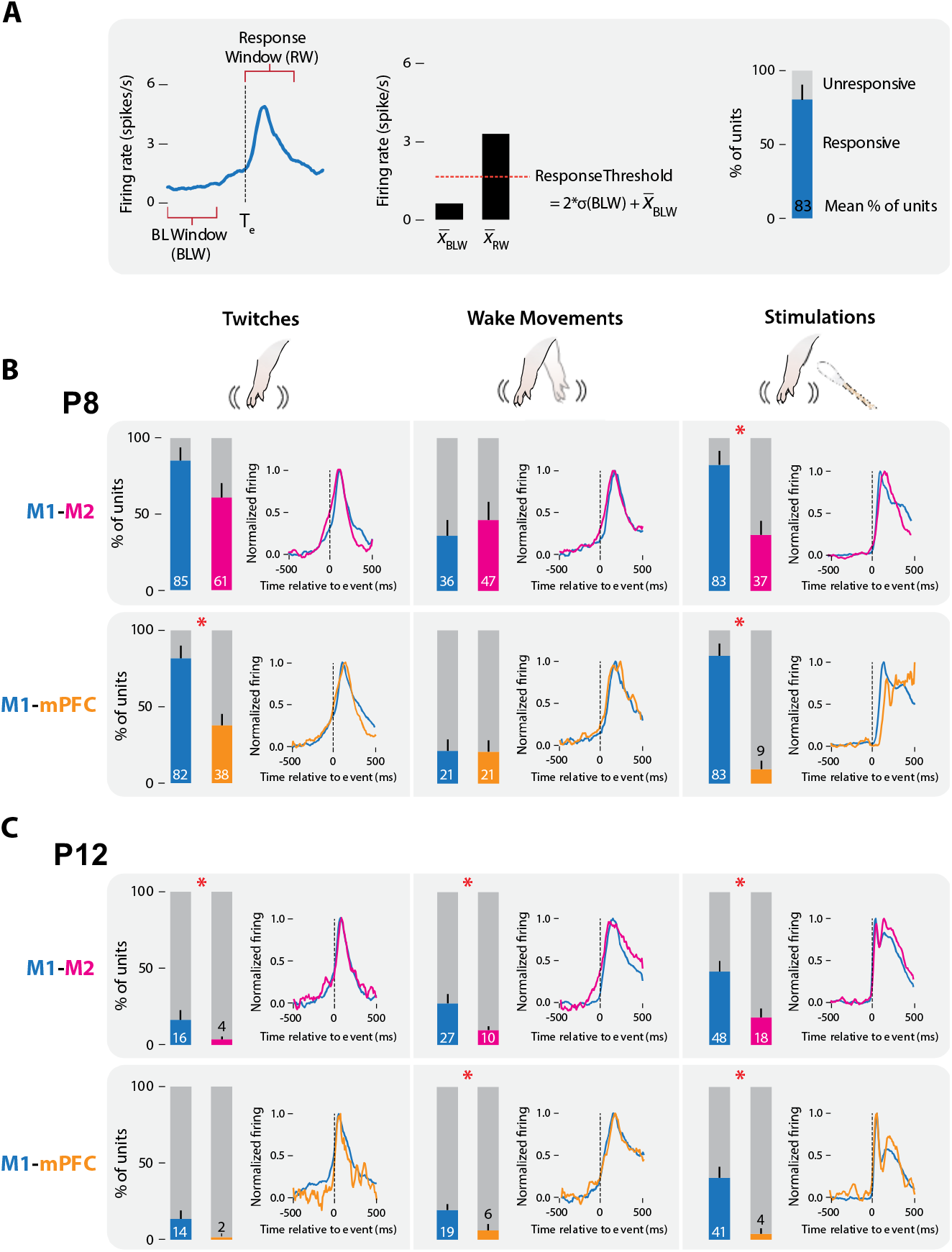
M1, M2, and mPFC neural responses to sensory input in P8 and P12 rats. (A) Methodology for determining sensory responsiveness of individual units. Left: Peri-event time histogram (PETH) of unit activity (blue line) surrounding a sensory event, showing the baseline window (BLW) and response window (RW). Event onset denoted by dotted line at T_e_. Middle: Bar graph showing mean unit activity during the BLW and RW, response threshold (dotted line), and the threshold calculation. Right: Stacked plot showing the percentage of units that exceeded the response threshold (blue) and units that did not (grey). (B) Stacked plots showing mean percentage of responsive (colored) and unresponsive (grey) units across pups at P8. Top: Data from M1 (blue) and M2 (magenta) recordings. Bottom: Data from M1 (blue) and mPFC (gold) recordings. Error bars are SEM. Asterisks denote significant difference between areas, p ≤ 0.017. PETHs to the right of each stacked plot show normalized activity profiles for responsive units only in each cortical area. (C) Same as in B, but at P12.

Regardless of the mean responsiveness of a cortical area at a given age, when units were responsive they exhibited response profiles (i.e., perievent time histograms; PETHs) that were strikingly similar to each other. These profiles indicate sensory processing as they include a clear peak response after the movement or stimulation (**Fig. 5B-C**). Such response profiles occurred even when responsive units were rare (e.g., mPFC at P12).

To assess whether M1, M2, and mPFC were similarly activated by sensory events, we next measured the activation rate. The activation rate was defined as the percentage of twitches, wake movements, or stimulations for which at least 30% of responsive units in the area showed an increase in activity (**Fig. 6A**). At P8 (**Fig. 6B**), M2 and mPFC had similar mean activation rates as those in M1 (t_(7)_s ≤ 2.28), with one exception: mPFC had a significantly lower mean activation rate than M1 for stimulations (t_(7)_ = 5.91, p < 0.001, Cohen’s D = 2.09). At P12 (**Fig. 6C**), mean activation rates in M2 and M1 were also similar (t_(8)_s ≤ 2.53); for mPFC, the mean activation rate for twitches was similar to that for M1 (t_(7)_ = 1.73), but mean rates were significantly lower in mPFC for wake movements and stimulations (t_(7)_s ≥ 4.06, ps ≤ 0.005, Cohen’s Ds ≥ 1.43). Finally, in contrast with previous findings in S1 and M1 (Gómez et al. 2021), we found no evidence that units in M1 are coactivated with units in M2 or mPFC (data not shown).

**Figure 6.**
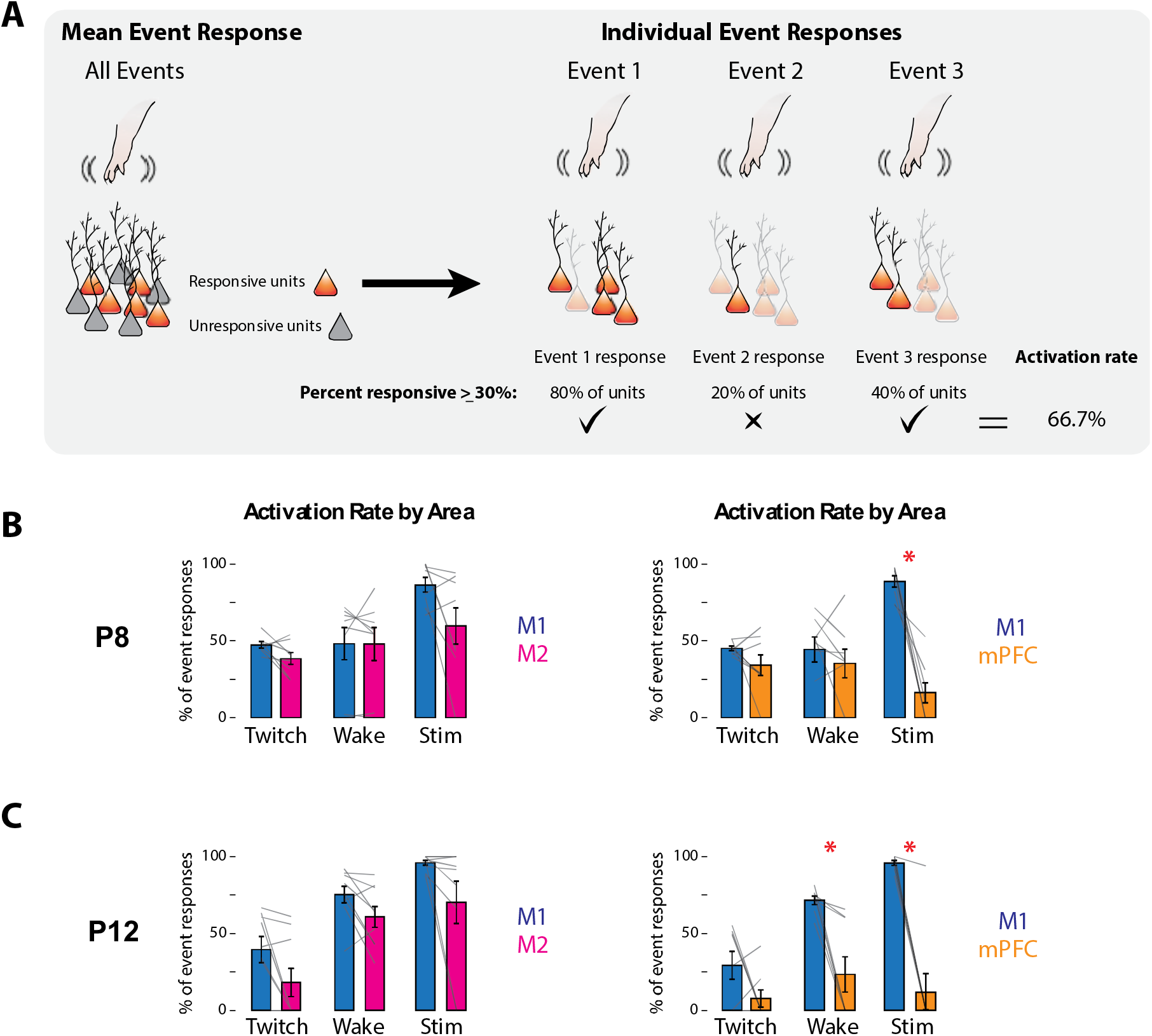
Response rates of M1, M2, and mPFC to sensory events in P8 and P12 rats. (A) Methodology for determining the responsiveness of cortical areas to sensory events. Left: Illustration of responsive (orange) and unresponsive (grey) units within an area. Right: Illustration of activity of responsive neurons (opaque) to individual sensory events. For each of the three sensory events indicated, the percentage of responsive units is determined. Based on the percentage of events that exceeds threshold (≥30%; check marks), the activation rate is calculated. (B) Activation rates in M1, M2, and mPFC at P8 to twitches (Twitch), wake movements (Wake), and stimulations (Stim). Left: Bar graphs showing percentage of sensory events that evoked a response in M1 and M2. Right: Same as at left, but for M1 and mPFC. Mean activation rates for individual pups are shown as grey lines. Error bars are SEM. Asterisk denotes significant difference between areas, p ≤ 0.017. (C) Same as in B, but at P12.

At P8 across all three areas, spindle bursts also reliably followed twitches, wake movements, and stimulations (**Fig. 6 – Supplement 1**). Congruent with unit responses, the mean probabilities of an event preceding a spindle burst were not significantly different between M1 and the other two areas, with the exception of twitches for M2 and stimulations for mPFC (t_(7)_s ≥ 3.51, ps ≤ 0.01, Cohen’s Ds ≥ 1.24). Thus, as shown previously in M1 and S1 at this age (Del Rio-Bermudez et al., 2020; Dooley et al., 2020; Glanz et al., 2021), sensory feedback triggers spindle bursts in M2 and mPFC.

In summary, similar to M1, M2 and mPFC respond to sensory input in early development with increases in spiking activity and spindle bursts.

### Urethane anesthesia suppresses behavior and neural activity in M1 and mPFC

The prefrontal activity described thus far does not resemble that reported in previous studies using urethanized pups (Brockmann et al., 2011; Bitzenhofer et al., 2015; Chini et al., 2019). To determine whether the use of urethane accounts for this disparity, we recorded M1 and mPFC activity in an additional set of P8 rats before and after administration of urethane (1.0 mg/g b.w. SC) or saline (n = 6/group).

Urethane administration produced rapid and dramatic effects on behavior and neural activity (**Fig. 7A**). Before urethane injection, pups cycled between sleep and wake and exhibited twitches and wake movements. In contrast, after urethane but not saline injection, limb movements decreased significantly (urethane: –87.14 ± 6.75%; t_(5)_ = 11.90, p < 0.001, Cohen’s D = 4.86; saline: +10.84 ± 5.84%, t_(5)_ = 1.00). The quality of limb movements also changed after urethane injection, with most movements consisting of brief whole-body spasms or rhythmic increases in muscle tone; twitch-like movements were rarely observed. These effects of urethane on limb movements and muscle tone precluded the identification of behavioral states.

**Figure 7.**
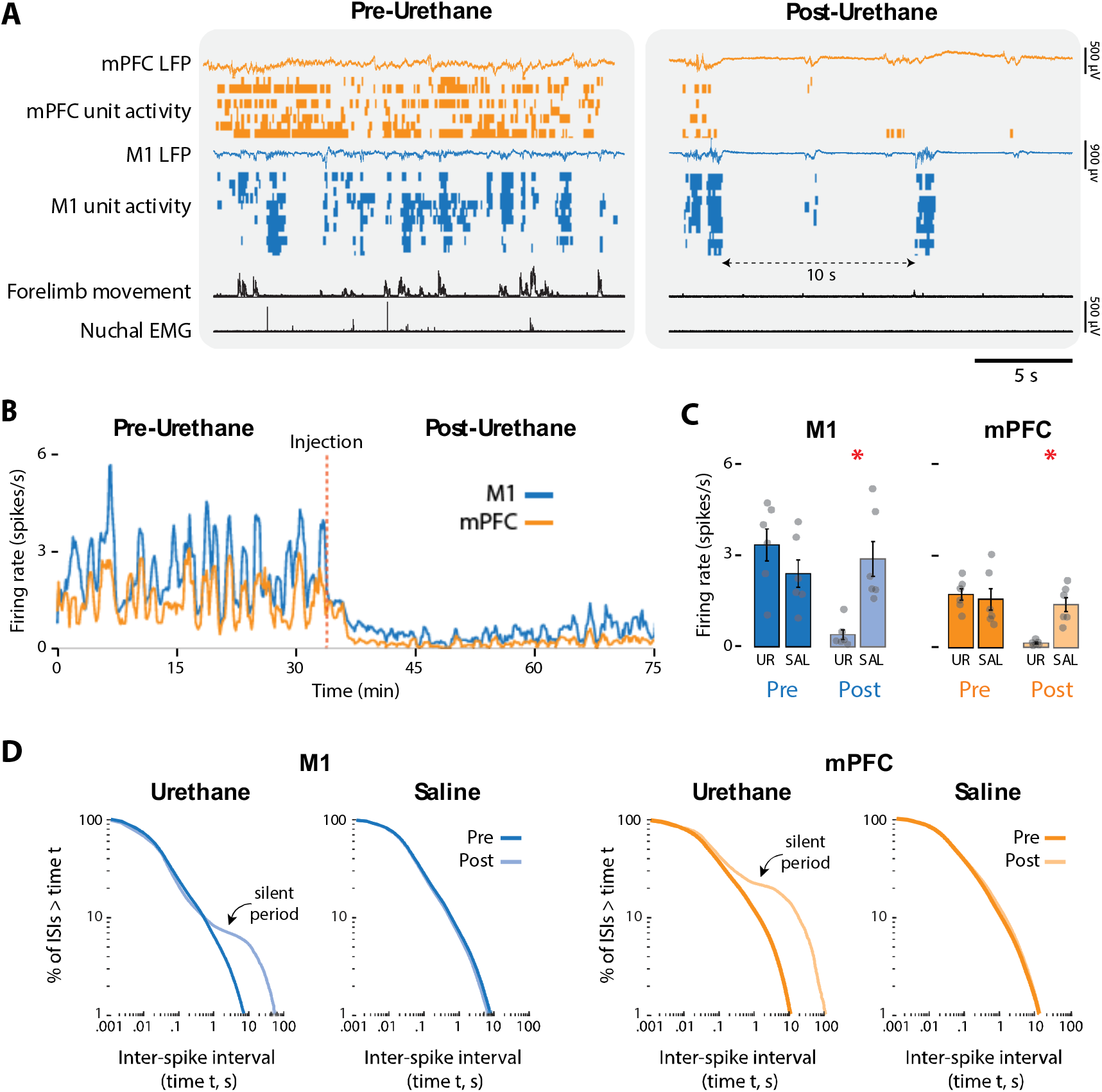
Urethane anesthesia suppresses unit activity in M1 and mPFC in P8 rats. (A) Representative 20-s segments of data from recordings in M1 and mPFC before (left) and after (right) injection of urethane (1.0 mg/g b.w.). For each record from the top, data are presented as follows: mPFC LFP (gold trace), mPFC unit activity (gold ticks), M1 LFP (blue trace), M1 unit activity (blue ticks), forelimb movement, and nuchal EMG. (B) Representative 75-min segment of data showing mean unit activity (2-s bins) in M1 and mPFC before and after injection of urethane (vertical dashed line). (C) Bar graphs showing mean firing rates of neurons across pups in M1 (left) and mPFC (right) during the pre-injection (Pre) and post-injection (Post) periods for the urethane (UR) and saline (SAL) groups. Mean firing rates for individual pups are shown as grey circles. Error bars are SEM. Asterisks denote significant difference between groups, p ≤ 0.025. (D) Left: Survivor plots of pooled inter-spike intervals (ISIs) for M1 units during the pre-injection (dark blue) and post-injection (light blue) periods for pups in the urethane and saline groups. Right: Same as at left but for mPFC during the pre-injection (dark gold) and post-injection (light gold) periods.

Urethane administration also disrupted neural activity in M1 and mPFC **(Fig. 7A-B**), causing reductions in firing rate of over 85% **(Fig. 7C**). Mean reductions in firing rate were significant for both M1 (t_(10)_ = 3.83, p = 0.003, Cohen’s D = 2.21) and mPFC (t_(10)_ = 5.01, p < 0.001, Cohen’s D = 2.94). Urethane also dramatically and significantly reduced the mean rate of spindle bursts in the two areas (t_(10)_s ≥ 3.18, ps ≤ 0.01, Cohen’s Ds ≥ 1.83) (**Fig. 7 – Supplement 1**).

Urethane also changed the spatiotemporal patterning of neural activity (**Fig. 7A**). In the absence of urethane, neural activity in both areas exhibited the discontinuous pattern characteristic of cortical activity at P8 (Golshani et al., 2009; Van Der Bourg et al., 2017; Glanz et al., 2021). In contrast, urethane injection produced a burst-suppression pattern that is characteristic of general anesthesia as well as coma, hypothermia, and neonatal trauma (Grigg-Damberger et al., 1989; Steriade et al., 1994; Hellström-Westas et al., 2006; Shanker et al., 2021), with intense population bursts separated by periods of relative silence commonly lasting 10 s or longer. These differences are illustrated by survivor plots of interspike intervals (ISI) (**Fig. 7D**): Whereas the pre- vs. post-injection ISI distributions for the Saline groups are indistinguishable, the post-injection distributions for the Urethane groups deviate substantially from the pre-injection distributions, due to the pronounced shoulders indicative of relative silence.

In summary, urethane anesthesia at P8 eradicates sleep-wake cycling, suppresses behavior, and produces an atypical pattern of neural activity.

## DISCUSSION

Here, we demonstrate in rats at P8 and P12 that neurons in two prefrontal areas—M2 and mPFC—exhibit state-dependent activity and responsivity to somatosensory stimuli. First, we find at both ages that neural activity in M2 and mPFC increases specifically during AS, similar to previous findings at these ages in M1 and S1 (Tiriac et al., 2014; Dooley et al., 2020; Glanz et al., 2021). Second, we find that neurons in M2 and mPFC respond to reafference arising from twitches and wake movements, and exafference arising from manual stimulation, with the proportion of responsive neurons generally being highest in M1 and decreasing across M2 and mPFC. Finally, we show that urethane thwarts accurate assessments of brain-behavior relations in developing cortex by suppressing neural activity and abolishing sleep-wake states, thus explaining discrepancies between the current and previous findings. Altogether, these findings highlight the potential importance of sleep and sensory experience for PFC’s functional development.

### Prefrontal cortex is most active during sleep

In developing rats, AS modulates spiking and oscillatory activity in S1 and M1 (Blumberg et al., 2020; Dooley et al., 2020; Glanz et al., 2021), findings that we now extend to M2 and mPFC. Neural activity in these two areas was highest during movement-related periods of AS, but it was also higher during AS than wake even in the absence of movement, thus suggesting state-dependent modulation. Accordingly, the present findings add to the growing body of evidence that state-dependent modulation is a general feature of developing cortex (Mukherjee et al., 2017; Blumberg et al., 2020; Del Rio-Bermudez and Blumberg, 2022).

In adults, state-dependent neuromodulation is a well-established phenomenon. For example, the basal forebrain provides AS-dependent cholinergic input throughout cortex (Lee & Dan 2012, Jones, 2020). However, little is known about the development and function of state-dependent neuromodulation in neonates. In prefrontal and visual cortex of newborn rats, acetylcholine from the basal forebrain modulates cortical responses (Hanganu et al., 2007; Janiesch et al., 2011). Likewise, monoaminergic neuromodulators, such as serotonin from the dorsal raphe, play formative roles in cortical development (Kolk and Rakic, 2022). However, in early development, the functional relations between neuromodulatory activity and the circuitry that governs sleep-wake states are unknown. Further research is needed to better understand state-dependent neuromodulation in developing cortex.

### Prefrontal cortex responds to sensory input

Sensory experience in early life is thought to scaffold the developing sensorimotor system, providing feedback about the growing body and the changing world it inhabits (Blumberg, 2015). In early development, M1 and S1 receive ample sensory input arising from self- and other-generated stimuli (Dooley and Blumberg, 2018; Glanz et al., 2021; Gómez et al., 2021). Sensory input to M1 and S1 is thought to refine somatotopy and connectivity within and between cortical areas (Dooley and Blumberg, 2018; Gómez et al., 2021) and, in M1, provide scaffolding for the development of motor functions. The conspicuous role for sensory experience in early development extends to higher-order areas such as M2, which has a coarse somatotopy related to its sensorimotor functions (Mohammed and Jain, 2014, 2016). The same cannot be said for mPFC, which does not exhibit somatotopic organization in either infants or adults. Indeed, it has been theorized that development of higher-order functions in prefrontal cortex derives in part from its relative independence from sensory input and reliance instead on intrinsic neural activity (Johnson et al., 2015, 2021; Werchan and Amso, 2017).

In light of the above, it was not surprising to find that units in M2, like those in M1, respond to sensory input from self- and other-generated movements of the forelimb. What was surprising, however, was the detection of sensory responses in mPFC. Indeed, responsive units in M1, M2, and mPFC exhibited nearly identical temporal profiles to sensory input, indicating similar modes of sensory processing. At the population level, the declining proportion of responsive units in the three areas—highest in M1 and lowest in mPFC—likely reflects their decreasing somatotopic homogeneity and increasing functional heterogeneity (**Fig. 8**) (Asanuma and Mackel, 1989; Bedwell et al., 2014; Barthas and Kwan, 2017). Thus, sensory input in early development likely serves different functional roles for each area.

**Figure 8.**
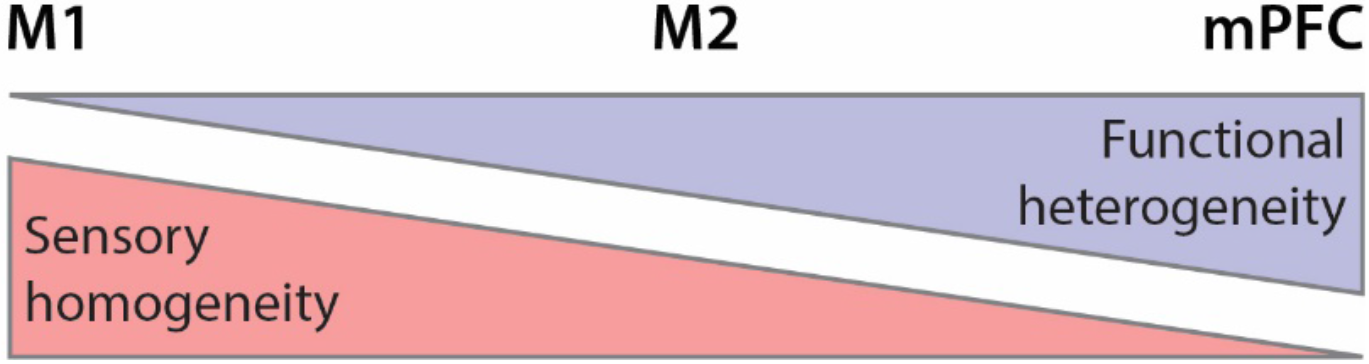
Summary illustration of sensory and functional gradients across M1, M2, and mPFC. Topographic organization of somatosensory input is more homogeneous in M1 than in M2 and mPFC (red). In contrast, functional organization is more heterogeneous in mPFC than in M2 and M1 (lavender).

Because the data presented in **Figure 4** are specific to the contralateral forelimb, our findings likely underestimate somatosensory responding in mPFC. Moreover, given that mPFC receives multimodal inputs (Hoover and Vertes, 2007; Bedwell et al., 2014), it is reasonable to expect that the developing mPFC also responds to olfactory, gustatory, visual, and/or auditory input. Indeed, in adults, mPFC responds to somatosensory and auditory input, which is thought to arrive via the posterior medial thalamic nucleus (POm, though this finding comes from heavily anesthetized subjects; see Martin-Cortecero and Nuñez, 2016). Thus, whereas unimodal sensory input to primary sensory areas enables the development of somatotopic homogeneity, multimodal sensory input to prefrontal cortex may enable the development of functional heterogeneity. To better understand the functional development of prefrontal cortex, future research should examine how multiple sensory modalities influence its early activity.

### Urethane abolishes brain-behavior relations

Until now, developmental investigations of rat prefrontal cortex were conducted in pups under urethane anesthesia (Brockmann et al., 2011; Bitzenhofer et al., 2015). Although urethane is known to alter neural activity and behavior (Dyer and Rigdon, 1987; Simons et al., 1992; Sorrenti et al., 2021), prior studies in rat pups have discounted the significance of these affects. In fact, it has been claimed that urethane mimics natural sleep in infant and adult rodents (Clement et al., 2008; Pagliardini et al., 2013; Bitzenhofer et al., 2015).

The present findings cannot be reconciled with these claims. Despite using a light dose, urethane rapidly suppressed neural activity and eradicated natural sleep-wake states and motor activity. In the absence of behavior, firing rates and oscillatory activity in both M1 and mPFC exhibited a pathological burst-suppression pattern of activity—activity that is rarely observed in healthy unanesthetized pups (Grigg-Damberger et al., 1989; Steriade et al., 1994; Hellström-Westas et al., 2006; Iyer et al., 2014; Shanker et al., 2021). Thus, as is increasingly appreciated in adults (Akeju and Brown, 2017; Mondino et al., 2022), anesthesia in infants is not a suitable proxy for natural sleep or sleep-related neural activity. Nor is it compatible with the goal of understanding brain-behavior relations in early development. Accordingly, extreme caution is warranted when reporting and interpreting results from anesthetized animals.

## Conclusions

Consistent with previous findings in sensorimotor cortex, we find that both behavioral states and sensory experience influence neural activity in developing PFC. These findings raise the possibility that the development of prefrontal cortex is influenced by more than intrinsic activity alone and that—similar to the once-unexpected sensory foundations of “motor cortex” in early development (Chakrabarty and Martin, 2005; Dooley and Blumberg, 2018; Glanz et al., 2021; Singleton et al., 2021)—the early expression of activity in PFC may not be related in obvious ways to the higher-order cognitive functions for which it is known in adults. Thus, our findings support the idea that state-dependent modulation and responsivity to sensory input are general features of developing cortex. In other words, it may be that the early functional development of primary and higher-order cortical areas are more similar than currently appreciated.

Finally, although the similarities and differences between rodent and primate PFC have been debated for decades (Preuss 1995; Carlén 2017; Barthas 2017; Laubach, 2018; Uylings 2003), there is still no consensus regarding the extent to which non-primates have cortical areas homologous to primate PFC. When defining PFC, different researchers variously emphasize anatomical connectivity and functional criteria, resulting in terminological confusion. For example, in rodents, M2 goes by many names and may perhaps be considered part of mPFC (Barthas 2017). The current study does not resolve these issues and was not designed to do so. But, by assessing sensory responses in M2 and mPFC in infant rats, this study introduces a developmental approach for comparing inter-areal PFC functionality across species. As has been shown for other cortical domains (Krubitzer and Dooley, 2013), a developmental-comparative approach holds the promise of clarifying PFC’s evolutionary and functional history.

## METHODS

### EXPERIMENTAL MODELS

All experiments were conducted in accordance with the National Institutes of Health Guide for the Care and Use of Laboratory Animals (NIH Publication No. 80–23) and were approved by the Institutional Animal Care and Use Committee of the University of Iowa.

Sprague-Dawley rats at P8–9 (hereafter “P8”; body weight: 20.76 ± 1.77 g) and P12–P13 (hereafter “P12”; body weight: 30.74 ± 2.16 g) were used. Pups were born to dams housed in standard laboratory cages (48 × 20 × 26 cm) with a 12-h light/dark cycle. Food and water were available ad libitum. The day of birth was considered P0 and litters were culled to 8 pups by P3. To protect against litter effects, pups selected from the same litter were always assigned to different experimental groups (Abbey & Howard 1973; Lazic & Essioux 2013).

#### Experiment 1: Recordings in unanesthetized subjects

In Experiment 1, we recorded M1, M2, and mPFC activity from unanesthetized pups at P8 and P12.

##### Experimental procedure

###### Surgical preparation

Surgery was performed using established methods (Blumberg et al., 2015; Glanz et al., 2021). Briefly, on the day of recording a pup of healthy weight and with a visible milk-band was removed from the litter, anesthetized with isoflurane (3.5–5%, Phoenix Pharmaceuticals, Burlingame, CA), and placed on a heating pad. Bipolar electrodes (California Fine Wire, Grover Beach, CA) were inserted into the nuchal, left, and right forelimb (*biceps brachii*) muscles and secured with collodion; an anti-inflammatory agent (Carprofen, 0.1 mg/kg SC; Putney, Portland, ME) was administered and the torso of the pup was wrapped in soft surgical tape. The scalp was sterilized with iodine and ethanol, and a portion of the scalp was removed to reveal the skull; a topical analgesic (bupivacaine, 0.25%; Pfizer, New Work, NY) was applied to the skull surface and surrounding skin, and then a veterinary adhesive (Vetbond; 3M, St. Paul, MN) was used to secure the skin to the skull. A steel head-fix (Neurotar, Helsinki, Finland) was attached to the skull using super glue (Loctite; Henkel Corporation, Westlake, OH) and dried with accelerant (INSTA-SET; Bob Smith Industries, Atascadero, CA). The pup was secured in a stereotaxic apparatus (Kopf Instruments, Tujunga, CA) where, under isoflurane anesthesia, a steel trephine (1.8 mm; Fine Science Tools, Foster City, CA) was used to drill openings in the skull over forelimb M1 (all coordinates from bregma; P8: +1.0 mm rostrocaudal (RC), 1.8 mm mediolateral (ML); P12: +1.0 mm RC, 1.8–2.0 mm ML), forelimb M2 (P8 and 12: +2.0 mm RC, 1.0 mm ML), and mPFC (P8 and P12: +1.8 mm RC, 0.5 mm ML). The pup was then transported to the recording rig, where it recovered for at least 1 h. Recording began only after regular sleep-wake cycles were observed and intracranial temperature reached 36°C.

###### Data acquisition

Neurophysiological and electromyographic (EMG) data were collected using a data acquisition system (Tucker-Davis Technologies, Gainsville, FL) with sampling rates of approximately 25 kHz and 1.5 kHz, respectively. Neural data were collected simultaneously from two locations using 16-site, 5-mm silicon-iridium electrodes (A1×16-3mm-100-177-A16 or A1×16-5mm-100-177-A16; NeuroNexus, Ann Arbor, MI). Before insertion, electrodes were coated with a fluorescent dye (DiI; Invitrogen, Waltham, MA) for later confirmation of placement. A chlorinated silver wire (0.25 mm in diameter; Medwire, Mt. Vernon, NY) was inserted into occipital cortex and used as both reference and ground. Neural data were recorded and visualized using Synapse software (Tucker-Davis Technologies). Video was collected using a BlackFly-S camera (100 fps) and SpinView software (FLIR Integrated Systems, Wilsonville, OR). To enable synchronization of the video and electrophysiological records, an LED light was positioned within the camera frame and was programmed to flash once every 3 s (Dooley et al., 2021).

###### Experimental design

We performed two sequential recordings in 10 pups at P8 (five female) and 14 pups at P12 (seven female). We first recorded activity from the forelimb regions of M1 and M2. Electrodes were inserted into the target sites and allowed to settle for at least 10 min. To confirm electrode placements in forelimb regions of M1 and M2 before recording began, the experimenter monitored the pup’s neural activity while using a cotton-tipped dowel to move the contralateral forelimb. On occasion, when forelimb-related activity was not detected during the first electrode placement, the electrode was withdrawn, repositioned, and lowered again; electrodes were never repositioned more than twice. Video and neurophysiological data were recorded for 40 min as the pup cycled freely between sleep and wake. This period was followed by 50 manual stimulations of the right forelimb (as described above), delivered approximately 2–3 s apart. Upon completion, the electrode in M2 was carefully withdrawn, coated again with DiI, and reinserted into mPFC (the M1 electrode was not disturbed). After the electrode settled in mPFC for at least 10 min, we again recorded video and neurophysiological data for 40 min, followed by 50 stimulations of the right forelimb. In total, the M1-M2 dataset consisted of eight recordings at P8 and nine at P12; the M1-mPFC dataset consisted of eight recordings at P8 and eight at P12.

##### Data analysis

###### Processing of neurophysiological data

Neurophysiological data were filtered for unit activity (bandpass: 300–5000 Hz) and converted into binary files. Templates for putative spikes were extracted using Kilosort (Pachitariu et al., 2016) and visualized using Phy2 (Rossant and Harris, 2013), as previously described (Dooley et al., 2021; Glanz et al., 2021; Gómez et al., 2021). Spike waveforms and autocorrelations were used to identify single units and multiunits. Preliminary analyses were performed to confirm that the activity profiles of single units and multiunits did not differ in any systematic way. Thus, all subsequent analyses were conducted using both single-unit and multiunit activity (hereafter “units” or “unit activity”). To obtain local field potentials (LFPs) neurophysiological data were down-sampled to approximately 1000 Hz, smoothed (0.005 s), and converted into binary files. Spike-time and LFP data were imported into MATLAB for analysis. To extract spindle bursts, LFP signals were bandpass filtered at 10–20 Hz (stopband attenuation: –60 dB; transition gap: 1 Hz) and the phase and amplitude of the filtered signal were calculated using a Hilbert transform (Glanz et al. 2021). Spindle bursts were defined as events for which the waveform amplitude exceeded, for at least 100 ms, the median amplitude plus 2x the standard deviation of the baseline amplitude. Spindle-burst onset was determined using previously described methods (Dooley et al. 2020).

###### Analysis of behavioral state

Motor activity and behavioral state were assessed visually using video and corroborated with EMG. We used custom-written MATLAB scripts to detect frame-by-frame changes in pixel intensity within user-defined regions of interest (Dooley et al., 2021). We selected regions encompassing the right forelimb and the rest of the body to allow detection of movement periods. Movements were represented as changes in pixel intensity across time. We then imported movement, neurophysiological, and EMG data into Spike2 (Cambridge Electronic Design, Cambridge, UK). Files were separated into periods of active sleep (AS) and wake (W); the remainder comprised periods of behavioral quiescence (periods that could not be clearly defined as sleep or wake) that were examined but not included in the present analyses. As defined previously (Del Rio Bermudez et al., 2020; Dooley et al., 2021; Glanz et al., 2021), AS is a period of nuchal muscle atonia accompanied by limb twitches (as identified from movement and EMG data); wake is a period of high nuchal muscle tone that is often accompanied by high-amplitude, coordinated limb movements. Scoring of behavioral state was performed blind to neural activity. To quantify differences in firing rate across AS and wake, for each unit we calculated the mean firing rate over the duration of each state; we then calculated mean firing rate across units within each brain area for each pup. Analyses were performed on these mean firing rates. At P8, we also assessed whether the occurrence of spindle bursts was state-dependent. For each area in each animal, we determined the mean rate of spindle bursts during AS and wake, and then calculated the mean spindle burst rate across pups. Spectrograms of oscillatory activity were generated using the sonogram function in Spike2. (This analysis was not performed at P12 because spindle bursts are not clearly discernable at this age.)

To delineate differences between state-dependent and movement-dependent changes in firing rate in M1, M2, and mPFC, we calculated the mean firing rate in each area during periods of movement. Movement and non-movement periods were extracted using custom MATLAB scripts from whole body movement data (derived from video as described above). The onset of a movement period occurred when movement data exceeded a threshold value of 3x greater than baseline for at least 250 ms; the offset of a movement period occurred when the movement data decreased below threshold. Movement and non-movement periods were categorized as to whether they occurred during AS or wake. Mean firing rates were calculated for each unit during the following conditions: AS with movement, AS with no movement, wake with movement, and wake with no movement. We determined the mean firing rate across units within an area and then the mean across animals.

###### Analysis of sensory activity

We analyzed neural activity in M1, M2, and mPFC in response to twitches, wake movements, and stimulations (hereafter referred to collectively as “sensory events”). First, we scored twitches of the right forelimb during AS using movement data and video. The onset of a twitch was defined as the first frame of movement. When a bout of rapid twitching was detected, the first twitch in the bout was always scored; subsequent twitches in the bout were also scored if they could be clearly distinguished from the previous twitch (e.g., by a change in movement direction). We also scored periods of wake for individual forelimb wake movements; as wake movements typically occurred as bouts of long continuous sequences, only the first limb movement in a bout was scored. The onset time of each forelimb stimulation was scored using video based on the first frame in which the dowel touched the forelimb. Scoring of sensory events was also performed blind to neural activity.

To determine whether units in M1, M2, and mPFC were responsive to sensory input, we constructed peri-event time histograms (PETHs) for unit activity triggered on sensory events. We calculated PETHs in spikes/s for each unit, triggered on sensory events (window = 1 s, offset = 0.5 s, bin size = 10 ms). We then defined a baseline window (BLW; -500 to -200 ms before the event) and a sensory-response window (RW; 0 to 200 ms after the event) and calculated the mean firing rate within each window. If the firing rate during the RW was greater than the mean baseline firing rate plus 2x the standard deviation of the baseline value 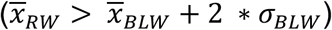, the unit was categorized as “responsive.” Using only responsive units, we again calculated average PETHs within each structure and normalized the data to the maximum value within each PETH.

To assess the reliability of response to a sensory event, we examined the activation rate, defined as the percentage of events to which each area responded. Again, using only responsive units, we examined the activity of each unit before and after each sensory event, using the same BLW as above but increasing the RW to 0– 300 ms after the event so that the window durations were equal. For a given unit within a cortical area, if the summed unit activity within the RW exceeded a threshold defined as 1.5x the summed unit activity within the BLW (∑ *RW* > 1.5 * ∑ *BLW*), the unit was categorized as having an event response. When more than 30% of recorded units within a cortical area exceeded this threshold, it was determined that the area as a whole for that pup was activated (Glanz et al. 2021). If a pup had no responsive units in a given area, that area’s activation rate was set to zero.

#### Experiment 2: Recordings in urethanized subjects

In Experiment 2, we used P8 rats to assess the effects of urethane anesthesia on behavior and neural activity in M1 and mPFC.

##### Experimental procedure

###### Surgical preparation

Surgical preparation. Pups were prepared for neurophysiological recording as in Experiment 1, with one exception: After securing EMG electrodes, a small (1 mm) incision was made in the skin near the base of the tail and surgical-grade silicon tubing (inner diameter: 0.020 in; outer diameter: 0.037 in; SAI Infusion Technologies, Lake Villa, IL) was secured subcutaneously with veterinary adhesive.

###### Data acquisition

Neurophysiological and video data were acquired as in Experiment 1.

###### Experimental design

We recorded from P8 rats before and after administering urethane (n = 7) or sterile saline (n = 6). We recorded baseline video and neurophysiological data for 30 min while pups cycled between sleep and wake, followed by 50 stimulations of the right forelimb as in Experiment 1. We then infused urethane (1.0 mg/g b.w.; Sigma-Aldrich, St. Louis, MO) or an equivalent volume of sterile saline (Fresenius Kabi, Bad Homburg, Germany) through the implanted cannula. This procedure minimized disruption of the pup and allowed for uninterrupted recording of data. Also, subcutaneous infusion of urethane produces a comparable level of surgical anesthesia as intraperitoneal injection and reduces the likelihood of organ puncture (Maggi and Meli, 1986; Field and Lang, 1988; Matsuura and Downie, 2000). We waited for at least 10 min for the drug to take effect, after which data were again recorded for 30 min followed by 50 stimulations of the right forelimb. One pup was excluded from analysis due to the complete loss of neural activity after urethane administration.

##### Data analysis

###### Processing of neurophysiological data

Neurophysiological and video data were processed as in Experiment 1; however, behavioral state could not be scored in this experiment due to the eradication of sleep-wake states by urethane.

###### Analysis of changes in movement quantity

To examine changes in the amount of movement before and after administration of urethane or saline, we quantified movement based on pixel-change data derived from video, as in Experiment 1. For each pup, we visualized smoothed (0.01 s) whole-body movement data in Spike2 to determine the baseline level of activity. Baseline movement activity was based on the mean pixel change across five 1-s windows during which no movement occurred. Then, in MATLAB, we summed the movement data (in pixel intensity changes per frame) separately during the pre- and post-injection periods; for each period, we performed a baseline subtraction. The sums of the movement data in the pre- and post-injection periods were divided by the total duration of each period. Next, we calculated the percentage change in movement between the pre- and post-injection periods. Finally, we calculated the mean percentage change across pups for each experimental group.

###### Analysis of changes in firing rate

We calculated the mean firing rate of each unit in M1 and mPFC during the pre- and post-injection periods. For each pup, we calculated mean unit firing rate within each area, and then calculated mean firing rates for each area across pups in the two experimental groups.

###### Analysis of changes in spindle-burst activity

We identified spindle bursts from LFP data as described in Experiment 1, using the pre-injection period to calculate the baseline LFP amplitude. The rate of spindle bursts was calculated for the pre- and post-injection periods.

##### Histology

At the end of each experiment, pups were overdosed with ketamine-xylazine (>0.08 mg/kg, IP) and perfused transcardially with phosphate-buffered saline (PBS, 1 M) followed by 4% paraformaldehyde (PFA). The brain was extracted and fixed for at least 24 h in PFA and 48 h in phosphate-buffered sucrose. Brain tissue was sliced coronally (80 µm) using a freezing microtome (Leica Biosystems, Wetlzar, Germany) and tissue was wet-mounted to locate the electrode tracks using fluorescence microscopy (2.5–5x magnification; Leica Microsystems). Sections were placed in well plates and stained for cytochrome oxidase (CO) to visualize cortical layers. Well plates were filled with CO solution (catalase, cytochrome C, DAB, phosphate buffered H2O, and DiH2O) and sections were allowed to develop in solution for 3-6 h on a heating pad (Dooley et al. 2021). Sections were then rinsed with PBS, slide-mounted, and allowed to dry for 48 h; slides were then placed in citrus clearing solution for 5 min, after which they were cover-slipped with DPX mounting medium. Cover-slipped slides were allowed to dry for at least 24 h. Fluorescent and brightfield images (at 2.5–5x magnification) were imported into Adobe Illustrator (San Jose, CA) and electrode tracks were reconstructed. CO-stained slides were used to determine the border between S1 and M1 (using layer 4 as a boundary). We demarcated mPFC and M2 based on the structure and orientation of cortical layers and with the aid of brain atlases (Paxinos and Watson, 2009; Khazipov et al., 2015).

### Statistical analyses

All statistical analyses were performed in SPSS (IBM) and MATLAB. For all tests, α was set to 0.05, unless otherwise specified; when appropriate, the Bonferroni procedure was used. A Shapiro-Wilk test was used to assess normality. We tested for significance using repeated-measures ANOVA and paired and unpaired t tests. Means are reported with their standard error (SEM). We report effect sizes for ANOVA as partial eta square (η_p_^2^), and for t tests as Cohen’s D.

## Data and Code Availability

Raw data (including timestamps of action potentials, oscillatory events, behavioral events, and behavioral states) will be uploaded to Dryad prior to publication. Select custom MATLAB scripts used here will up uploaded to Github prior to publication (https:\\www.github.com\XXXXX).

## Acknowledgments

This research was supported by grants from the National Institutes of Health (R37-HD081168 to MSB). We thank Greta Sokoloff, Ryan Glanz, and J. Toby Mordkoff for helpful comments and advice.

## Author Contributions

Conceptualization, LJG, JCD, and MSB; Methodology, LJG, JCD, and MSB; Software: LJG; Formal Analysis: LJG; Investigation: LJG; Data Curation: LJG; Writing – Original Draft, LJG; Writing – Review & Editing, LJG, JCD, and MSB; Visualization: LJG, JCD, and MSB; Funding Acquisition, MSB; Resources, MSB; Supervision, JCD and MSB.

---

**Figure 6 – Supplement 1.**
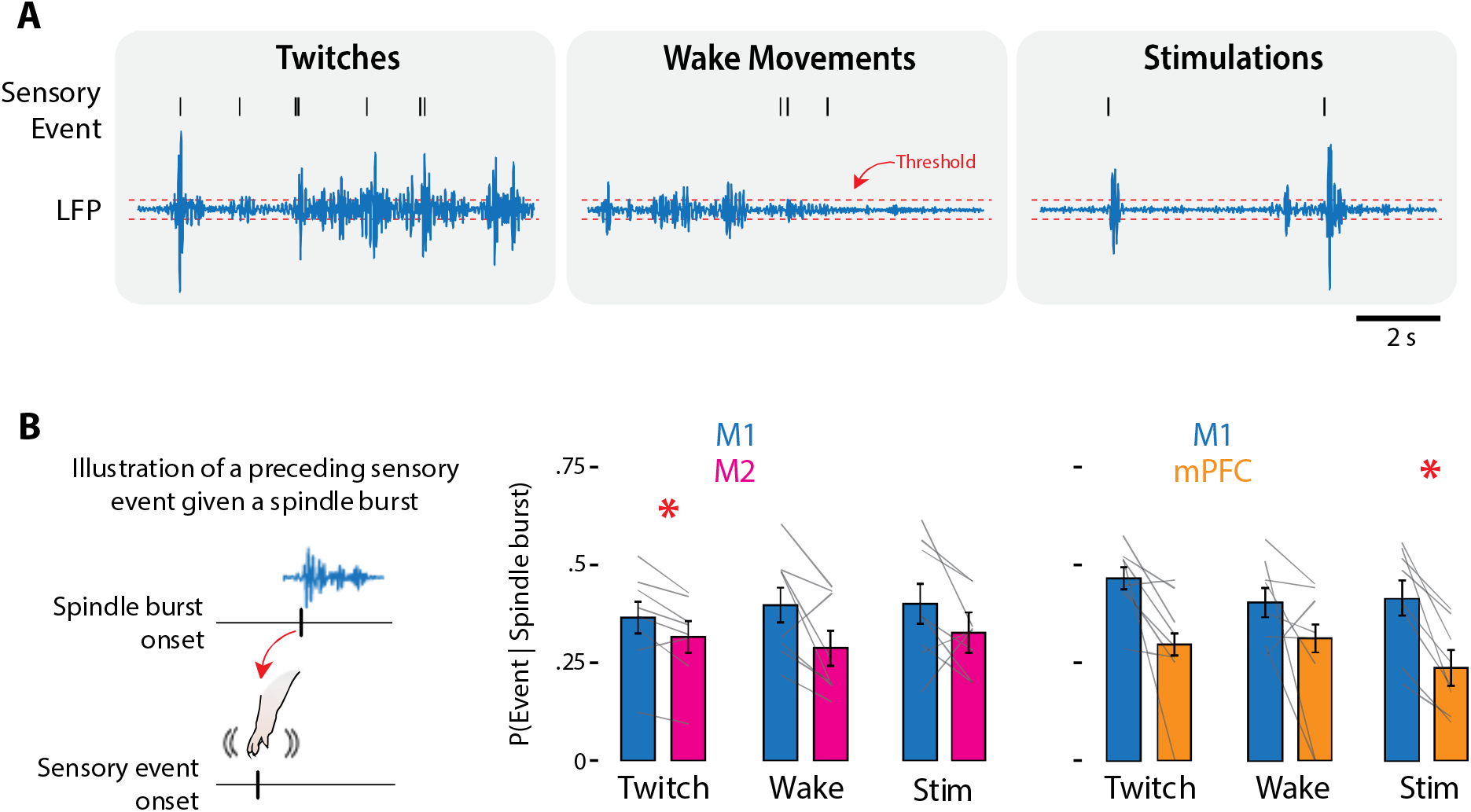
Spindle bursts in M1, M2, and mPFC in response to twitches, wake movements, and stimulations in P8 rats. (A) Filtered LFP (10–20 Hz) during periods of twitching, wake movements, and stimulations. (B) Left: Illustration of a spindle burst given a preceding sensory event; Right: Bar graphs showing the probability of a spindle burst given a preceding sensory event for M1, M2, and mPFC to twitches (Twitch), wake movements (Wake), and stimulations (Stim). Mean probabilities for individual pups are shown as grey lines. Error bars are SEM. Asterisks denote significant difference between areas, p ≤ 0.017.

**Figure 7 – Supplement 1.**
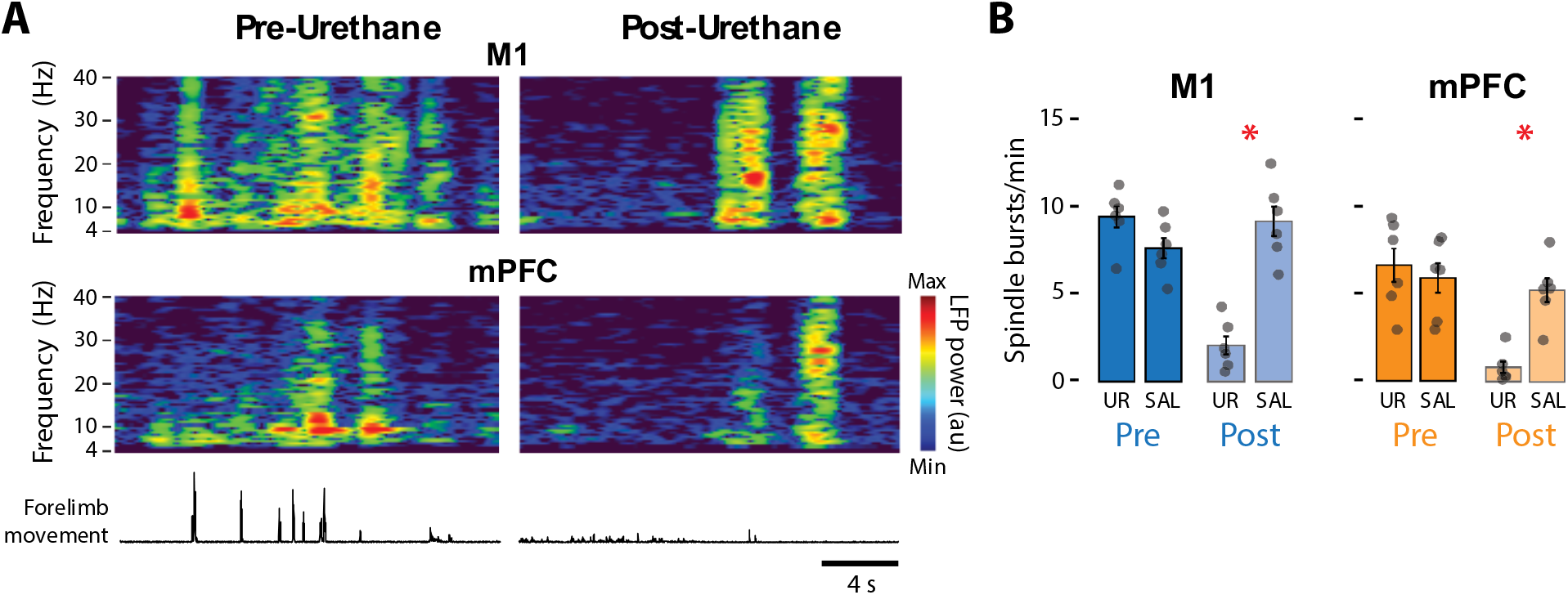
Urethane anesthesia reduces the rate of spindle bursts in M1 and mPFC in P8 rats. (A) Representative spectrograms showing 20-s of data pre- and post-urethane administration in M1 (top) and mPFC (bottom). Forelimb movements are shown below. (B) Bar graphs showing mean spindle-burst rate across pups in M1 (left) and mPFC (right) during the pre-injection (Pre) and post-injection (Post) periods in the Urethane (UR) and Saline (SAL) groups. Mean spindle-burst rates for individual pups are shown as grey circles. Error bars are SEM. Asterisks denote significant difference between groups, p ≤ 0.025.

